# DEET as a feeding deterrent

**DOI:** 10.1101/174979

**Authors:** WeiYu Lu, Justin Hwang, Fangfang Zeng, Walter S. Leal

## Abstract

The insect repellent *N,N*-diethyl-3-methylbenzamide (DEET), is a multimodal compound that acts as a spatial repellent as well as an irritant (contact repellent), thus being perceived by the insect’s olfactory and gustatory systems as an odorant and a tastant, respectively. Soon after DEET was developed, almost 6 decades ago, it was reported that it reduced mosquito feeding on blood mixed with this repellent. It is now known that the mosquito proboscis senses contact repellents with the tips (labella) of the labium, which remain in direct contact with the outer layers of the skin, while the stylets, including the feeding deterrent sensor (labrum), penetrate the skin. We designed a behavioral assay that allowed us to tease apart contact repellency from feeding deterrence. First, we demonstrate here that when DEET was mixed with blood and covered by Parafilm® layers, it did not leak to the outer surface. In our assays, the mean number of landings and duration of contacts with surfaces covering blood mixed with DEET or blood plus solvent (dimethyl sulfoxide) did not differ significantly. The feeding times, however, were significantly different. When blood was mixed either with 0.1 or 1% DEET, female southern house mosquitoes spent significantly less time feeding than the time spent feeding on blood mixed only with the solvent. By contrast, there were no significant differences in the mean times of feeding on blood containing 1% picaridin and blood plus solvent. Like DEET, the contact repellent and insecticide, permethrin, caused a significant reduction in feeding time. We, therefore, concluded, that in this context, DEET and permethrin act as feeding deterrents.

## Introduction

Chemicals used to reduce mosquito bites are not only repellents sensu stricto, ie, compounds that cause the responder to steer away from the source, but are also excitorepellents or irritants, ie, chemicals eliciting increased locomotor activity after an insect makes contact with the source (Obermayr 2015). From a strict mechanistic viewpoint, these 2 groups should be named noncontact and contact disengagents, respectively (Miller et al. 2009). From a more pragmatic perspective, the end result is the same, ie, mosquitoes are kept at bay by sensing odorants in the vapor phase (spatial repellents) and/or by detecting non-volatile tastants (contact repellents) upon direct contact with these chemicals (on a skin surface, for example). Although its complete mode of action is still a matter of considerable debate, DEET (=*N,N*-diethyl-3-methylbenzamide) is undoubtedly a multimodal compound (DeGennaro 2015), which is perceived by both the olfactory and gustatory systems as an odorant and a tastant, respectively. Additionally, evidence in the literature suggests that DEET also acts as a feeding deterrent (Barzeev & Smith 1959). The pioneering findings by Bar-Zeek and Schmidt (Barzeev & Smith 1959) that blood-feeding was prevented when samples were spiked with DEET has been overlooked most probably because of the difficulty in teasing apart feeding deterrence from contact repellency.

Mosquitoes sense the environment with their antennae, maxillary palps, proboscis, tarsi, and ovipositors. Whereas the antennae and maxillary palps are involved in the reception of odorants (eg, spatial repellents), the proboscis is involved in the reception of contact repellents and other tastants. This sophisticated “microneedle system” (Kong & Wu 2010) comprises a gutter-like labium that encloses a fascicle. There are 2 lobes (labella) at the tip of the labium, and the fascicle contains 6 stylets: a pair of teeth-bearing maxillae, a pair of mandibles, a hypopharynx with its salivary canal, and a labrum that carries sense organs on its tip (Wahid et al. 2003). During feeding, the fascicle penetrates the host’s skin while the labium bends and the labella remain in direct contact with the outer layer of the skin (Choo et al. 2015). Although it has been demonstrated that labral apical sensilla respond to phagostimulants (Liscia et al. 1999; Werner-Reiss et al. 1999) and feeding deterrents (Kessler et al. 2014), it remains difficult to unambiguously determine whether reduced feeding on DEET-spiked blood is mediated by “contact repellency” or “deterrence.” Indeed, Bar-Zeek and Schmidt (Barzeev & Smith 1959) suggested that “repellency” was caused by low concentrations of DEET (then named diethyltoluamide) in the blood.

To address whether reduced feeding on DEET-spiked blood was due to repellency or deterrence, we devised a modified version of our surface landing and feeding assay (Fig. 1) (Leal et al. 2017). We lured mosquitoes to feed on 2 cotton rolls covered with dual layers of Parafilm® sealing film and loaded with blood, one spiked with DEET and the other with solvent, and meticulously measured feeding times in the 2 parts of the arena. Here, we report that mosquitoes spend significantly less time feeding on DEET-spiked blood than on the control. Likewise, permethrin also acted as a feeding deterrent, but the time spent feeding on blood spiked with picaridin was not significantly different from the time spent on feeding on the control side of the arena.

**Figure 1.**
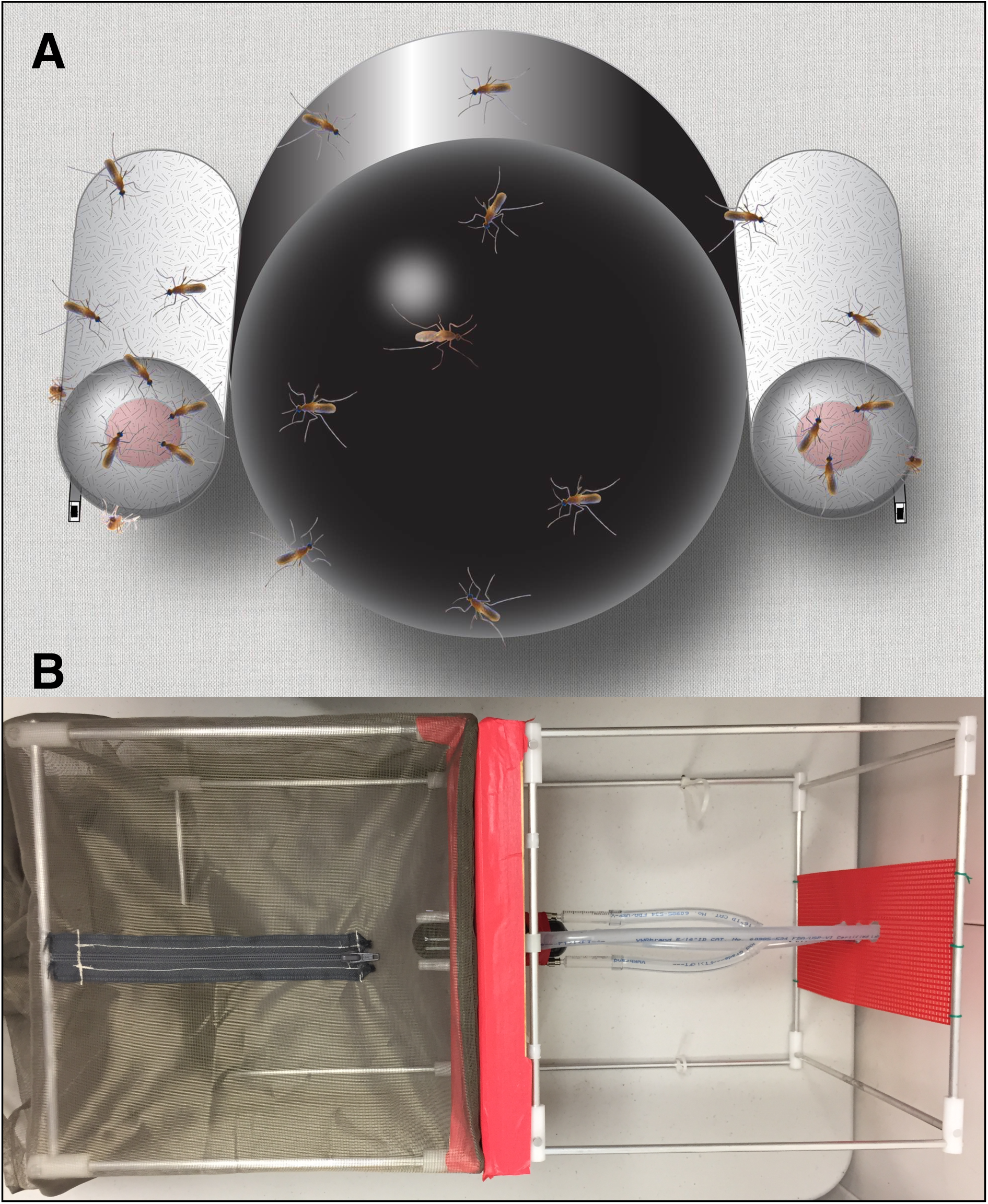
Illustration of the modified arena. (**A**) A Dudley tube painted black from inside was flanked by 2 cotton rolls secured in place by syringe needles that delivered CO_2_. Samples of defibrinated sheep blood mixed with solvent only or spiked with DEET were loaded on these cotton rolls, which were subsequently covered with Parafilm. (**B**) An aerial view of the arena. Mosquitoes were placed on a mosquito cage accessible from the top and having a camera (not shown) attached to the left. The Dudley tube was connected to a water bath (not shown) and the syringe needles to a CO_2_ tank (not shown).

## Materials and methods

### Mosquitoes

*Culex quinquefasciatus* mosquitoes used in this study were originally from a laboratory colony initiated with mosquitoes collected in the 1950s in Merced, California and currently kept by Dr. Anthony Cornel (Kearney Agricultural Center, University of California-Davis). The Davis colony has been maintained separately for more than 6 years under 12:12 (L:D), 27±1°C, and 75% relative humidity.

### Behavioral arena

Feeding behavior was measured using a modified surface landing and feeding assay (Leal et al. 2017). In brief, the device consisted of a base and a detachable assay cage (Fig. 1B). The frame of the base was made from an aluminum collapsible field cage (Bioquip, 30.5 × 30.5 × 30.5 cm) with a wooden board (30 × 30 cm) attached to the front of the cage and covered with red cardstock (The Country Porch, GX-CF-1) and red lab tape. Three openings were drilled through the wooden board to accommodate one 50-mL Dudley bubbling tube (Fisherbrand, 40356) and two 16-gauge syringe needles (Sigma-Aldrich, Z108782), orientations of which are illustrated on Fig. 1A. The Dudley tube painted internally with black hobby and craft enamel (Krylon, SCB-028) was attached to a water bath circulator with the temperature set at 38°C. The 2 syringe needles were connected to a CO_2_ tank through a bubbler to deliver CO_2_ at 50 mL/min. The frame of the detachable assay cage was made with the same aluminum collapsible field cage. Red cardstock was taped internally at 1 face of the cage, 1 circular opening, and 2 small holes were made in the cardstock to allow the Dudley tube and CO_2_ needles to project into the mosquito cage. The cage was completed with a field cage cover (Bioquip, 30.5 × 30.5 × 76.2 cm). One square, sealable opening (7 × 7 cm) was made at the backside of the field cage cover, allowing the Dudley tube and CO_2_ needles to insert into the cage. A slit was made on the top of the cage, and a zipper (10 cm) was sewn on to the slit for an easily accessible opening. A camera-accessible opening (d=5 cm) with a drawstring was made at the front of the field cage (Fig. 1B).

### Chemicals

DEET and permethrin (mixture of cis and trans isomers) were acquired from Sigma-Aldrich (PESTANAL^®^, analytical standards); picaridin was a gift from Dr. Kamal Chauhan (USDA-ARS, Beltsville) (Leal et al. 2017). Stock solutions (10% m/v) were prepared in dimethyl sulfoxide (DMSO) and diluted to 1% when needed. The blood mixtures were prepared by mixing 180 µL of defibrinated sheep blood (UCD, VetMed) with 20 µL of a 10% solution (of DEET, picaridin, or permethrin) to give a final concentration of 1%. The control was prepared in the same manner but using only DMSO.

### Behavioral measurements

Fifty female mosquitoes (6 days after emergence) were aspirated and transferred to the arena 2 hours before each experiment. All openings were sealed, and the cage was kept near the base of the arena. Thirty minutes after the water started circulating, the assay cage was then inserted into the base (Fig. 1). Aliquots (200 µL) of blood mixed with DMSO only or DEET in DMSO were gently pipetted onto one end of a piece of dental cotton (Primo Dental Products, #2 Medium) to make a blood circle on the cotton. A strip of Parafilm sealing film (ca. 8 x 5 cm) was stretched fully along the length and then wrapped around the cotton roll, covering the surface twice. To distinguish the treatment from the control group, a snipped insect pin (BioQuip, black enameled No.5) was tagged at the back of the cotton by a small piece of Parafilm. The sealed cotton rolls were placed in between the CO_2_ dispensing needles and the Dudley tube. Five microliters (the amount of 1 blood meal (Nikbakhtzadeh et al. 2016)) of defibrinated sheep blood were smeared onto the surface of the Parafilm (to prime mosquitoes to start feeding). CO_2_ flow was initiated, and the assay was recorded with a camcorder equipped with a Super NightShot Plus infrared system (Sony Digital Handycan, DCR-DVD 910). After 30 min, insects were gently removed from the cotton rolls, and the assays were reinitiated with fresh sealed cotton rolls with switched positions. For each group of tested mosquitoes, test and control were placed at least twice on each side of the arena.

### Statistical analysis

Behavioral observations were not done in real time, but rather by retrieving the recorded videos. Mosquito-feeding duration was counted only after the blood used for priming was already dried. For measuring feeding time, we selected mosquitoes that clearly pierced the membrane by forcing its head down towards blood, stopped movement of the head and the body, and started waving the hind leg while the stylets were inserted. Once all these steps were observed, we rewound the tape and started counting the feeding time. End of feeding was determined when the proboscis was removed and mosquitoes walked away. We preferred mosquitoes that were feeding solitarily rather than in groups so as to avoid interruption of feeding by other mosquitoes’ interference. We limited observations to at most 10 mosquitoes per assay, but each experiment was replicated 3-9 times and comparisons were made at least 30 times. Treatments and their controls were compared by 2-tailed Wilcoxon matched-pairs signed rank tests using Prism 7 (GraphPad, La Jolla, CA).

## Results and discussion

### Behavioral responses

Upon retrieving the videos, it became clear that contact repellency was not involved. Indeed, the mean duration of landings on the treatment side of the arena did not differ significantly (Wilcoxon 2-tailed, matched-pairs signed rank test, n=3, P<0.05) from the mean duration of landings on the control side (Fig. 2A). Additionally, the mean time that mosquitoes spent on the Parafilm-covered blood spiked with DEET did not differ from the mean time spent on the surface covering blood devoid of DEET (Fig. 2B). Of note, this “residence time” on the Parafilm surfaces was recorded from the time mosquitoes landed and before feeding was initiated. As far as contact is concerned, mosquitoes behaved similarly when landing on the surfaces covering blood spiked with DEET or loaded with blood plus solvent. These observations suggest that DEET did not leak from the blood to the outer surface of the paraffin film. Therefore, the feeding times we measured next were not influenced by repellency upon contact with the surfaces. We observed that mosquitoes probed similarly on both sides of the arena; the difference in behavior was observed once they had initiated a blood meal (Video 1). Mosquitoes spent significantly more time (91.8±12.1 s) feeding on the control side of the arena than on cotton rolls loaded with 0.1% DEET-spiked blood (32.7±4.2 s, n=30; P<0.0001, Prism notation: ****) (Fig. 3A). Likewise, they spent significantly less time feeding on 1% DEET-spiked blood (30.8±2.1 s, n=90) than on blood with solvent only (78.6±8.2 s, n=90; P<0.0001, ****) (Fig. 3B). Surprisingly, there was no significant difference in the time feeding on blood spiked with 1% picaridin (76.6±11.2 s) compared with its control (89.0±7.2 s, n = 60; P=0.0364) (Fig. 3C). Although all samples were freshly prepared and tested, we cannot rule out the possibility that picaridin degraded more rapidly upon being mixed with blood.

**Figure 2.**
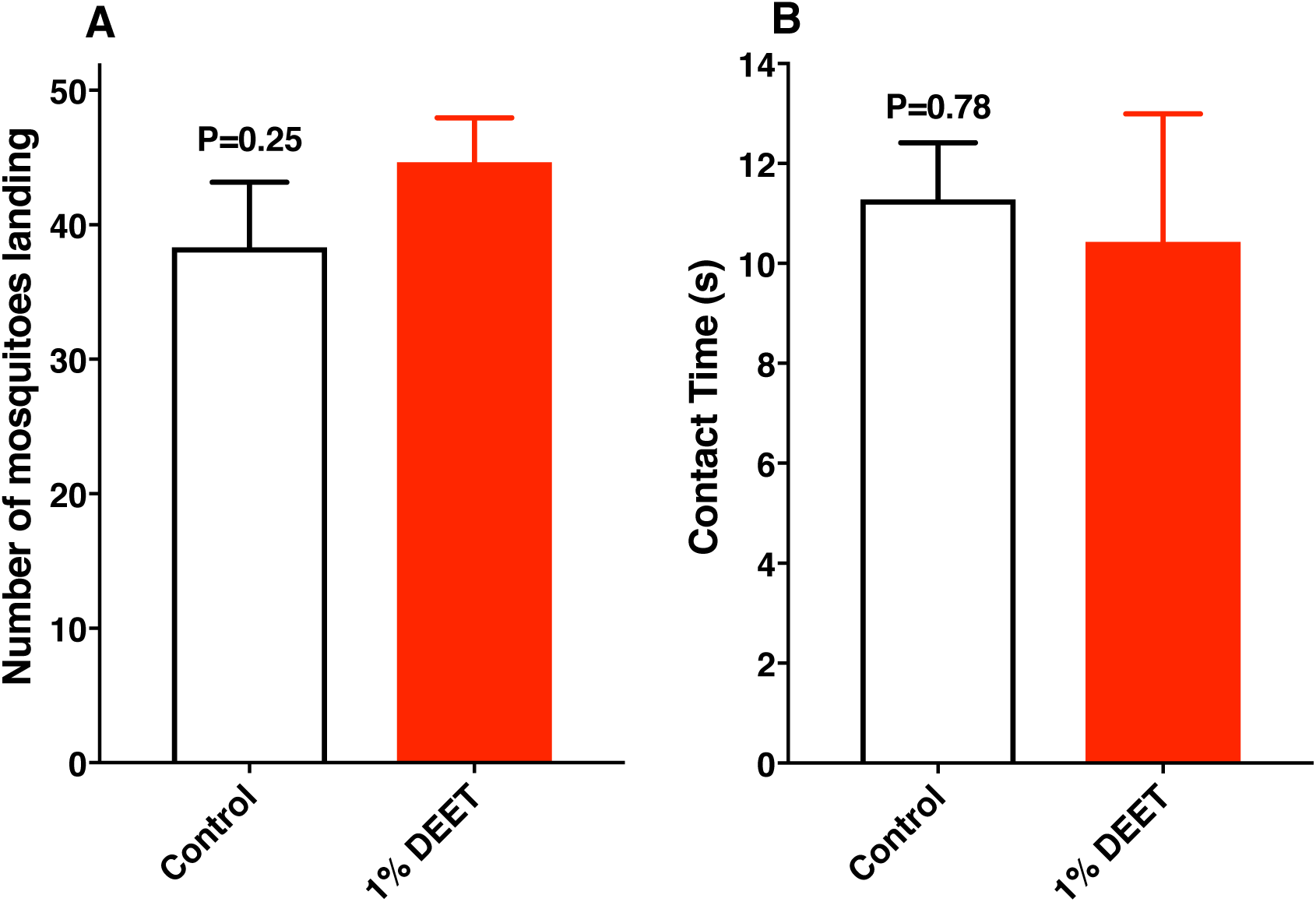
Measurements of landings and duration of contact with the surfaces prior to feeding. (**A**) The mean number of mosquitoes landing on the control and DEET sides of the arena in 15 min did not differ significantly (Wilcoxon matched-pairs signed rank test, n=3). (**B**) The contact times measured from the time the mosquitoes landed until they started feeding were not significantly different (Wilcoxon matched-pairs signed rank test, n=7).

It has been demonstrated that a DEET-sensitive odorant receptor from the southern house mosquito, CquiOR136, (Xu et al. 2014) is also expressed in the tip of the labrum (Choo et al. 2015). Therefore, we initially surmised that mosquitoes detected DEET in the blood samples by activating this receptor. The fact that this receptor is sensitive to both DEET and picaridin coupled with the lack of feeding deterrence elicited by picaridin does not support this assumption. It is, therefore, likely that mosquitoes detect DEET in the blood with their gustatory system. Next, we tested the effect of permethrin, a compound commonly used in long-lasting insecticidal nets (Kawada et al. 2014) given its dual property as an insecticide and excitorepellent (Zaim et al. 2000). Of note, permethrin is neither a spatial repellent nor a ligand for CquiOR136 (Xu et al. 2014). Like DEET, permethrin had a significant deterrent effect, with mosquitoes feeding significantly less on permethrin-spiked blood (21.8±2.8 s) than on blood containing only DMSO (79.6±8.8 s, n= 60; P<0.0001, ****) (Fig. 3D).

**Figure 3.**
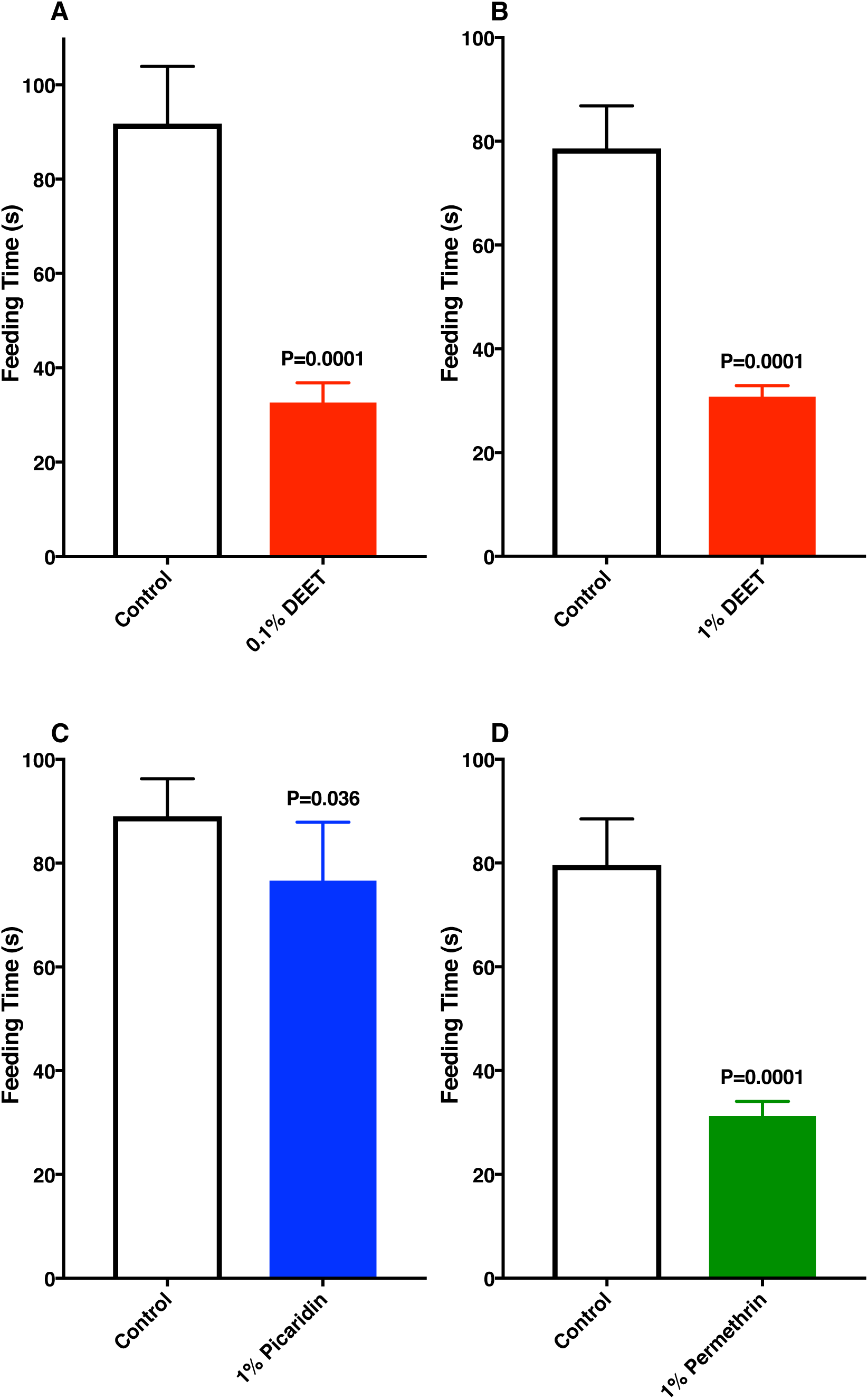
Comparative feeding times on blood mixed with solvent or test repellents. (**A**) 0.1% DEET, (**B)** 1% DEET, (**C**) 1% picaridin, and (**D**) 1% permethrin.

## Conclusions

With a modified version of the surface landing and feeding assay (Leal et al. 2017), we were able to demonstrate that reduced feeding on blood spiked with DEET was due to a deterrent rather than contact repellency effect. In this experimental setup, we provided blood on cotton rolls, which were covered with 2 layers of Parafilm. DEET did not leak and, consequently, contact repellency was not at play. This is demonstrated by the fact that mosquitoes landed randomly on the various surfaces of the arena (Video 1) and that the number and duration of the landings on the surface covering blood spiked with DEET did not differ from the similar data recorded for the side covering blood with solvent only (Fig. 2). Upon direct contact of the stylets with blood, mosquitoes prematurely terminated feeding on blood spiked with DEET and permethrin, but not with picaridin. Our findings suggest that the earlier observation of “repellency” by the presence of DEET (Barzeev & Smith 1959) in blood is due to “feeding deterrence.” In addition to being a spatial and a contact repellent, DEET is also a feeding deterrent. Previously, it has been suggested that DEET is a feeding deterrent due to contacts with treated surfaces (Klun et al. 2006). By contrast, our findings show that feeding is deterred by direct contact with a blood meal. Whereas the 2 well-known properties of DEET are essential for reducing mosquito bites and, consequently, transmission of diseases, “feeding deterrence” is of less importance in medical entomology given that once mosquitoes are already in contact with the blood they may have already transmitted arbovirus.

## Additional Information and Declarations

### Competing interests

No author has competing interests to disclose.

### Author contributions

WSL designed the experiments and constructed the behavioral arena. WL, JH, and FZ carried out the research. WL, JH, FZ, and WSL analyzed the data. WSL and WL wrote the manuscript. All authors have agreed to the final content of the manuscript.

## Data Availability

All raw data are provided as Supplementary Information.

### Funding

This work was supported in part by the National Institute of Allergy and Infectious Diseases of the National Institutes of Health under awards R01AI095514 and R21AI128931. FZ was supported in part by the Chinese Scholarship Council.

## Acknowledgements

We thank Dr. Anthony “Anton” J. Cornel (University of California, Department of Entomology & Nematology) for providing mosquitoes that allowed us to duplicate his colony at the Davis campus and Dr. Kamal Chauhan (USDA, ARS, Beltsville) for providing a sample of picaridin used in this research.

## References

Barzeev M, and Smith CN. 1959. Action of Repellents on Mosquitoes Feeding through Treated Membranes or on Treated Blood. J Econ Entomol 52:263–267.

Choo YM, Buss GK, Tan K, and Leal WS. 2015. Multitasking roles of mosquito labrum in oviposition and blood feeding. Front Physiol 6:306. 10.3389/fphys.2015.00306

DeGennaro M. 2015. The mysterious multi-modal repellency of DEET. Fly (Austin) 9:45–51. 10.1080/19336934.2015.1079360

Kawada H, Ohashi K, Dida GO, Sonye G, Njenga SM, Mwandawiro C, and Minakawa N. 2014. Insecticidal and repellent activities of pyrethroids to the three major pyrethroid-resistant malaria vectors in western Kenya. Parasit Vectors 7:208. 10.1186/1756-3305-7-208

Kessler S, Gonzalez J, Vlimant M, Glauser G, and Guerin PM. 2014. Quinine and artesunate inhibit feeding in the African malaria mosquito Anopheles gambiae: the role of gustatory organs within the mouthparts. Physiological Entomology 39:172–182. 10.1111/phen.12061

Klun JA, Khrimian A, and Debboun M. 2006. Repellent and deterrent effects of SS220, Picaridin, and Deet suppress human blood feeding by Aedes aegypti, Anopheles stephensi, and Phlebotomus papatasi. J Med Entomol 43:34–39.

Kong XQ, and Wu CW. 2010. Mosquito proboscis: An elegant biomicroelectromechanical system. Phys Rev E 82: 011910.

Leal WS, Barbosa RM, Zeng F, Faierstein GB, Tan K, Paiva MH, Guedes DR, Crespo MM, and Ayres CF. 2017. Does Zika virus infection affect mosquito response to repellents? Sci Rep 7:42826. 10.1038/srep42826

Liscia A, Crnjar R, Barbarossa IT, Esu S, Muroni P, and Galun R. 1999. Sensitivity of the mosquito *Aedes aegypti* (Culicidae) labral apical chemoreceptors to phagostimulants. J Insect Phys 39:261–265.

Miller JR, Siegert PY, Amimo FA, and Walker ED. 2009. Designation of chemicals in terms of the locomotor responses they elicit from insects: an update of Dethier et al. (1960). J Econ Entomol 102:2056–2060.

Nikbakhtzadeh MR, Buss GK, and Leal WS. 2016. Toxic Effect of Blood Feeding in Male Mosquitoes. Front Physiol 7:4. 10.3389/fphys.2016.00004

Obermayr U. 2015. Excitorepellency. In: Debboun M, Frances SP, and Strickman DA, eds. Inset Repellents Handbook. Boca Raton, FL: CRC Press, 91–115.

Wahid I, Sunahara T, and Mogi M. 2003. Maxillae and mandibles of male mosquitoes and female autogenous mosquitoes (Diptera: Culicidae). J Med Entomol 40:150–158.

Werner-Reiss U, Galun R, Crnjar R, and Liscia A. 1999. Sensitivity of the mosquito Aedes aegypti (Culicidae) labral apical chemoreceptors to phagostimulants. J Insect Physiol 45:629–636.

Xu P, Choo YM, De La Rosa A, and Leal WS. 2014. Mosquito odorant receptor for DEET and methyl jasmonate. Proc Natl Acad Sci U S A 111:16592–16597. 10.1073/pnas.1417244111

Zaim M, Aitio A, and Nakashima N. 2000. Safety of pyrethroid-treated mosquito nets. Med Vet Entomol 14:1–5.

